# Towards a universal structural and energetic model for prokaryotic promoters

**DOI:** 10.1101/348854

**Authors:** A. Mishra, P. Siwach, P. Misra, B. Jayaram, M. Bansal, W.K. Olson, K.M. Thayer, D.L. Beveridge

## Abstract

With almost no consensus promoter sequence in prokaryotes, recruitment of RNA polymerase (RNAP) to precise transcriptional start sites (TSSs) has remained an unsolved puzzle. Uncovering the underlying mechanism is critical for understanding the principle of gene regulation. We attempted to search the hidden code in ~16500 promoters, of twelve prokaryotes representing two kingdoms, in their structure and energetics. Twenty eight fundamental parameters of DNA structure including backbone angles, base pair axis, inter base pair and intra base pair parameters were used and information was extracted from X-ray crystallography (XRC) data. Three parameters (solvation energy, hydrogen bond energy and stacking energy) were selected for creating energetics profiles using in-house programs. DNA was found to be inherently designed to undergo a change in every parameter undertaken, from some distance upstream of TSSs to adopt a signature state at these locations in all prokaryotes. These signature states might be the universal hidden codes recognised by RNAP. This observation was reiterated when randomly selected promoter sequences (with little sequence conservation) were subjected to structure generation; all developed into very similar three dimensional structures, quite distinct from those of conventional B-DNA and coding sequences. Fine structural details at important motifs (viz. −11, −35, −75 positions relative to TSS) of promoters reveal novel and pointed insights for RNAP interaction at these locations; it could be correlated that how some particular structural changes at −11 region may allow insertion of RNAP amino acids in inter-base pair space as well as facilitate the flipping out of bases from DNA duplex.

## Introduction

An organism’s complete set of genetic information is expressed in a highly regulated manner across time and space. Sequence elements within or near core promoter regions of genes contribute to regulation (1,2), but there is almost no universal consensus promoter sequence in prokaryotes (3,4). Recently discovered non-canonical transcripts in prokaryotes also have unconventional promoter location and architecture, as revealed by genome-wide transcriptional start sites (TSSs) mapping studies at single nucleotide resolution (5). What guides the recruitment of transcriptional machinery so precisely to so many unconventional sites? Structural homology among different promoters, where different sequences lead to similar structural variants, was considered as an alternative criterion quite early^3^. Lately, DNA structural descriptors like DNA stability, stacking energy, A-philicity, propeller twist, roll among others have been used to define/identify promoter regions, to a certain extent, in both prokaryotes and eukaryotes and some structural properties were found to correlate well with gene expression (6–10). Though these studies make a significant contribution towards understanding of promoter architecture, things are far away from a universal model capable of explaining the underlying mechanism of transcription initiation at precise locations. RNA polymerase is considered as the central component in transcription regulation, regulating by recognizing and binding to specific promoter sequences and facilitating unwinding of DNA duplex near TSS. With emerging reports on DNA structure regulating biological processes (11), a need arises to know-whether promoter structure acts simply as a passive platform on which transcriptional machinery (RNA polymerase and sigma factors in bacteria) acts or it also regulates/directs/actively participates in transcription initiation.

The study was planned with two clear goals: to prepare complete structural and energetic profiles of TSSs and their adjoining regions in search for a universal model for prokaryotic promoter and to understand their implications on transcription initiation. The structure and dynamics of nucleic acids is guided by base sequence as well as by the sugar-phosphate backbone. Earlier attempts, mentioned above, have focussed only on sequence and that too by taking only a few parameters; no attempts have been made towards complete structural and energetic characterization of promoter regions. Last few decades have witnessed a revolutionary evolution in the analysis of nucleic acids structure (12–20). We have previously reported that hydrogen bond, stacking and solvation energy show clear signatures of functional densities of DNA sequences (21–27).

For the present study, we proceeded with nine backbone, eight inter-base pair (inter-BP), six intra-base pair (intra-BP), five base pair-axis (BP-axis), and three energetic properties adding to a total of thirty one parameters for exploring the genomic regions comprising primary TSSs, of twelve microorganisms (belonging to both kingdoms-archea and eubacteria, of prokaryotes). Numeric values of conformational parameters for the unique di-nucleotides steps were obtained from crystal structures of B-DNA only (from nucleic acid database (NDB) using curves+ (17), while in-house programs were used for energy parameters (27). Here, we report that these parameters provide unique structural and energetic signatures at TSSs. Our results offer fundamentally new insights into the active role of DNA structure and energetics at TSS in transcription initiation and offer new pathways to explore transcriptional regulation in prokaryotes.

## Materials and methods

### Promoter and coding sequence dataset preparation

A total of 16519 primary TSS positions were selected from twelve organisms (Table 1). Sequences of 1001 nucleotides length (spanning 500 nucleotides upstream and downstream of the TSS positioned at 0), for all selected TSSs positions were extracted from respective genome sequence. As control dataset, coding sequence (CDS) data for the respective organism were retrieved from Ensemble bacteria site. Out of 45,220 CDS sequences, only 6218 sequences had length greater than 1500nt, from which we extracted 1001 central region as control dataset for our analysis.

### Crystal structures of B-DNA only

A total of 74 crystal structures of B-DNA, without any modification or association with protein or ligand molecule, were obtained from NDB database (see supplementary Table S1).

### Structural parameter value calculation

Twenty eight parameters were selected-nine backbone (Alpha, Beta, Gamma, Delta, Epsilon, Zeta, Chi, Phase and Amplitude), eight inter-BP (Shift, Slide, Rise, Tilt, Roll, Twist, H-Rise and H-Twist), six intra-BP (Shear, Stretch, Stagger, Buckle, Propel and Opening), five BP-axis (X Displacement, Y Displacement, Inclination, Tip and Axis-Bend. The values for these parameters, for the crystal structures obtained above, were calculated using Curves+ (17). After calculating values for all the parameter for each B-DNA structure, all occurrences of unique ten dinucleotide steps in the 5’ to 3’ direction were considered for each parameter and the average of all the occurrences was calculated. The parameter values for the unique dinucleotide steps thus obtained are provided in supplementary Table S2.

### Energy parameter value calculation

The values for three energy parameters viz. Hydrogen bond energy, stacking energy and solvation energy for the unique ten dinucleotide steps was done as reported in our previous work (27).

### Obtaining the structural and energy profile of each sequence

The calculated dinucleotide values for each parameter were used for getting structural profile of 1001 nucleotide long promoter and CDS sequence by performing moving average calculation on sliding window of 25 base pair covering 24 dinucleotide steps. The same exercise was performed independently on all the selected sequences of primary promoter sequence and CDS sequence (as control) for all the 31 parameters.

### Profile plotting of sequences

The plotting was performed using MATLAB software.

### Normalization of values

To bring all the parameters on the same sacle, the values were made dimensionless using normalization. The values were normalized between 0 and 1 by subtracting the minimum value of the profile from each value and then divide the value with range of the profile (i.e. max - min).

### Making derived structural criteria to define a sequence

The normalized values, showing similar behaviour were combined together to form two structural vectors; vector1 from 14 parameters showing peak (Stretch, Opening, Rise, Roll, Twist, H-Rise, H-Twist, Beta, Gamma, Epsilon, Phase, Amplitude, Hydrogen bond, Stacking energy) while vector2 from 17 parameters showing cleft at TSS (X Disp., Y Disp., Inclination, Tip, Ax-Bend, Shear, Stagger, Buckle, Propel, Shift, Slide, Tilt, Alpha, Delta, Zeta, Chi, Solvation).

### Generating Structures of promoter DNA

Twelve sequences (−75 to +25) were extracted with respect to randomly selected TSSs, one from each organism and were subjected to structure generation using X3DNA software package (28). Fine structures of five nucleotide long motifs (from −11, −35 and −70 regions) of *Bacillus amyloliquefaciens* were also generated. We first generated generic B-DNA structure of selected sequences using fiber tool of the 3DNA package and then analyzed the structures with the help of find_pair and analyze tool. This command generated two parameter values files, base pair step parameter file and base pair helical parameter file. In the first step, we modified the base pair step parameter value file using our predicted value and generated modified pdb structure using rebuild tool. Then we again analyzed the modified pdb structure using find_pair and analyze tool. This time, we modified the base pair step helical parameter file using our predicted values and rebuild the second step modified pdb structure. In this way, we are able to modify values for 18 DNA structural parameters including inter-BP, intra-BP and BP-axis parameters i.e. all except the backbone angles and sugar puckering variables. Since all parameters are correlated, it is assumed that these 18 structural parameters are sufficient to generate the structure of DNA sequence.

## Results and Discussions

### All structural and energy parameters give signature profiles at Transcription start sites

Primary promoter sequences were obtained by extracting five hundred nucleotides both upstream and downstream to the given TSS from the complete genome sequence of each organism while coding sequences (CDSs) were obtained from Ensemble Bacteria site and only the central region (one thousand nucleotides long) of each CDS was taken (Table 1).

**Table 1:**
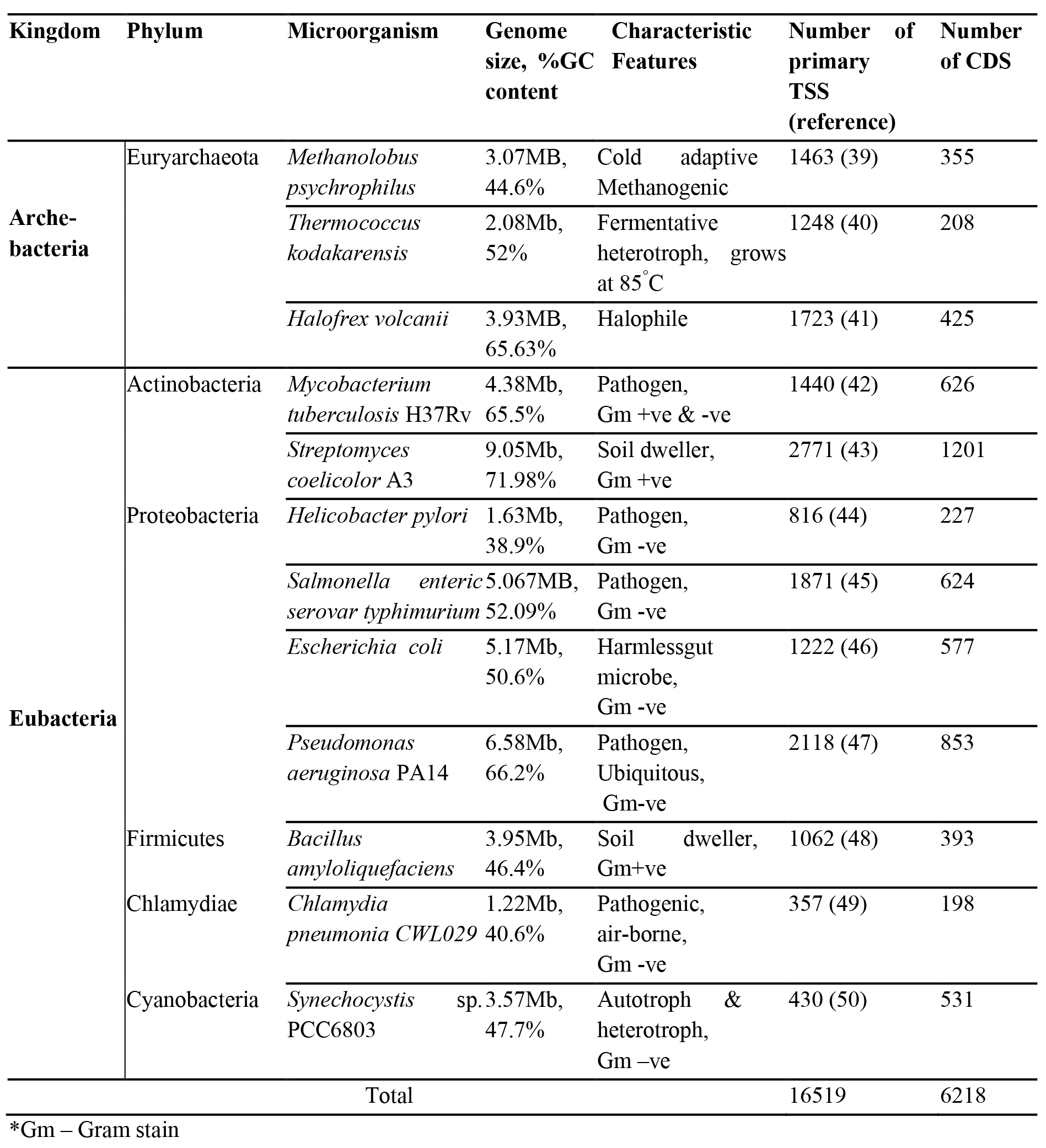
A brief description of the selected microorganisms along with the transcriptional start sites and coding sequences data, used in the present study.

Numeric profiles of thirty one structural and energy parameters were obtained for the pooled primary promoters (16519) and the CDSs (6218) (see Methods) and are shown in Fig. 1.

**Fig 1.**
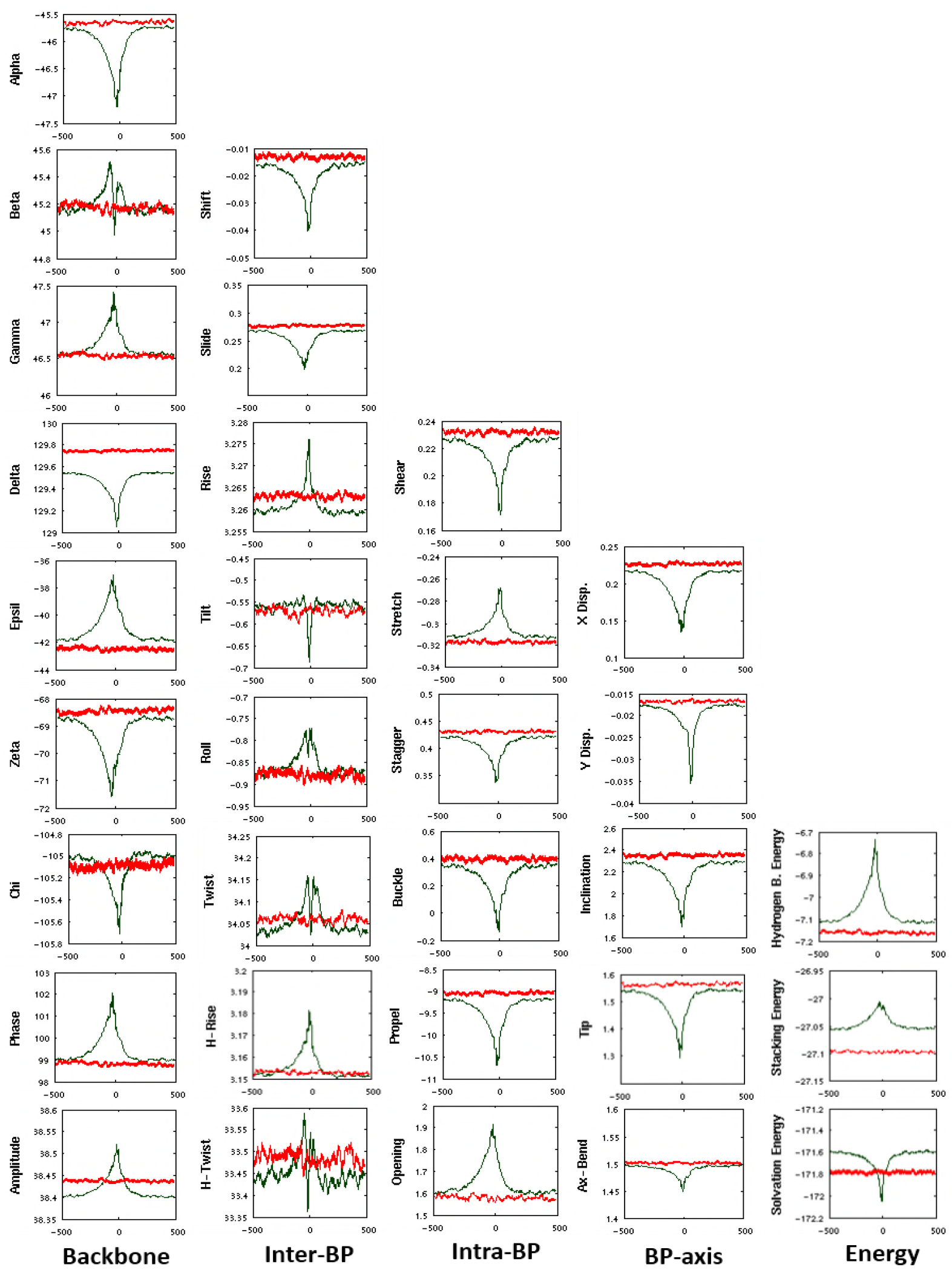
Structural and energy profiles of 1001 nucleotides long sequences having primary promoters (green line) and coding sequences (red line). Sequences, having primary promoters, (16519) were lined up with TSS at the same position (“0”), extending 500 nucleotides on both sides. Likewise all CDSs were also superimposed The ordinate represents the numeric value of that parameter, while abscissa represents the nucleotide position.

These pooled profiles were obtained by lining up all the promoter sequences with TSS at the same position, and all CDSs were also superimposed. Next, all the sequences were converted to numeric sequences for different structural parameters, and the average over all numeric sequences for each position is plotted (Fig. 1). The abscissa shows the position relative to TSS while the ordinate represents the numeric value of that parameter. As clear from Fig. 1, all the parameters are capable of distinguishing primary promoter sequences from CDSs. The promoter sequences show unique intrinsic value at TSS and nearby regions resulting in a sharp/broad peak/cleft at/near transcriptional start sites (TSSs), and hence make a signature profile for that parameter (Fig. 1; for individual profiles of each organism see Supplementary Fig. SI).

As the sequence proceeds to TSS, a gradual increase in the basal value (given by CDS and extreme upstream and downstream regions of TSS) is observed for thirteen parameters (beta (β), gamma (γ), epsilon (ε), phase, amplitude, rise, H-rise, roll, twist and H-twist, stretch, opening, solvation energy) while eighteen properties (alpha (α), delta (δ), zeta (ζ), chi (χ), shift, slide, tilt, shear, stagger, buckle, propeller twist, x-displacement (Xdis), y-displacement (Ydis), inclination, tip, ax-bend, hydrogen bond energy, stacking energy) show gradual decrease, till TSS or its nearby upstream position and afterwards re-track back to basal values. Correlation exists among these structural properties but ultimately each parameter contributes in its own way. The results obtained were analyzed to fulfil this need to know the impact of each parameter on the overall structure and shape of DNA at TSS, as presented below.

Some properties adopt a very gradual change pattern spanning across a long distance (from −250^th^±100 position through TSS to +100^th^±50) almost in all the twelve prokaryotes. This category includes twenty four properties-all torsion angles (α, β, γ, δ, ε, ζ, χ) and sugar puckering variables (phase and amplitude) of sugar phosphate backbone, all the five BP-axis parameters (Xdis, Ydis, inclination, tip and ax-bend), six inter-BP parameters (shift, slide, roll, H-rise), four intra-BP properties (shear, stretch, stagger, buckle) and two energy properties (hydrogen-bond and stacking energy (Fig.1; Supplementary Fig. SI). The second category belongs to those properties which give very sharp signature profile, spanning across a small length of 30 to 35 nucleotides or less (−20±5 to +10±5); it includes seven parameters-four inter-BP (rise, tilt, twist, H-twist), two intra-BP (propeller twist and opening) and one physicochemical property (solvation energy) (Fig.1; Supplementary Fig. S1). Either the required change, in each of these properties, at TSS can be achieved by following their respective pattern or the change itself is needed across the respective distances.

The B-DNA backbone is realized in two major conformer sub-states: BI and BII, inter-conversion guided by coupled changes in two dihedral angles ε, ζ. BI sub-state is characterized by lower value of ε and higher values of ζ (with ε-ζ<0), while reverse (with ε-ζ>0) is true for BII sub-state (29). Though the values of torsion angles ε and ζ, observed in the present study do not coincide with that of the canonical B-DNA, but their dynamics, as the sequence proceed towards TSS, correlate with transition from BI to BII sub state (ε increases while ζ decreases) and at TSS ε attains maximum while ζ has the minimum value (i.e. backbone appears to be in BII conformer) in all prokaryotes (Fig. 1; Supplementary Fig. SI). BII is the less common sub state of B-DNA as has been observed in crystal structures and molecular dynamics simulations (30). Another way to define the backbone transitions is to look at α, γ angles which are found to associate with canonical and non-canonical backbone states; with α decreasing while γ increasing during transition from canonical to non-canonical state (31). The similar negative coupling between α, γ angles was observed as the sequences proceed to TSS (with α decreasing and γ increasing) in all the selected prokaryotes; indicating a trend from canonical state to a non-canonical state, though the angles values were far from standard values given for canonical/non canonical (Fig. 1).

Base pairs of promoters show an increasing tendency to align on top of each other, as the sequences move towards TSS, by gradually decreasing shift and slide values. However, the increased angular distance between base pairs towards minor groove side (i.e. roll) does not allow the base pairs to be in parallel. Roll dynamics exhibits some peculiar trend: while undergoing a gradual increase it shows a sudden decrease near −35^th^±10 position followed by sudden rise and then a slow decrease till past TSS (Fig 1). A similar trend was also observed for twist and H-twist except that the sudden decrease followed by sudden increase was observed near −10^th^±5 position. Rise increases while tilt decreases across almost same span (−20±5 to +10±5) (Fig. 1). The inter-BP parameters, obtained from atomic molecular dynamics simulations, have been used earlier in promoter prediction algorithm (8).

Among the various intra-BP parameters, a gradual decrease is observed for shear, buckle, and stagger resulting in centrally aligned bases on the intersection of x and y-axis. The base pairs show a gradual increase in stretch (from ~−250^th^±100bp), with peak near −10^th^±5 followed by gradual decrease. Propeller twist shows a sharp decrease (making the base pairs to be more parallel to y-axis) while opening shows a sharp increase at around −10^th^±5 position. Propeller twist has also been reported earlier as a differentiating property between promoters and non-promoters (6,7).

BP-axis of promoter regions was observed to have lower values of translational (Xdis, Ydis) and rotational movements (tip, inclination) as well as of Ax-bend, compared to adjoining regions. Decrease in Xdis and Ydis move the base pairs towards centre along x-axis and y-axis. Likewise decrease in rotational movements (inclination and tip) would orient the base pairs to adopt perpendicular orientation to axis. It can be said that the helix becomes narrow and rigid, and bases more perpendicular to the axis, as the sequences proceed to TSS. Less bendability of promoter regions around TSS has also been reported earlier (9, 10).

Among the three physicochemical properties, hydrogen bond energy and stacking energy exhibit a gradual decrease when the sequence moves towards TSS till around −10^th^±5 position, afterwards showing a gradual increase till past TSS. Sharp increase in solvation energy was observed at around −10^th^±5 position of the promoter sequence. Lesser stability of promoter region has also been reported earlier (10, 26).

At individual prokaryote level, it is observed that prokaryotes differ greatly in the mean genomic value and signal strength at TSS for a given parameter, but the nature of change is almost similar (Supplementary Fig. SI). Further, the difference in the mean genomic value and the signal strength at TSS for a given parameter is not found to correlate with genome size, phylogeny and %GC content.

### Combining all parameters for obtaining a single criterion

These parameters were made statistically unit-less so as to evaluate on a single scale (see Methods). When these thirty one normalized (dimensionless) parameters of all the twelve organisms (31 × 12) are plotted together on this new structural scale, a clear peak and cleft is observed at TSS or its adjoining upstream region (Fig. 2).

Next step was to join together all the parameters so as to make a single structural criterion to define local DNA structure. As discussed in the previous section, some properties show a gradual increase while others a gradual decrease till TSS, combining all together would nullify each other’s effect and will result in reduced signal. So instead of a single vector, two structural vectors were made by joining together all the parameters with similar behaviour: vector1 from parameters showing peak while vector2 by combining parameters showing cleft (see Methods). When values of these two vectors were plotted for the promoters and CDSs, a three line graph was obtained for all organisms-single line for CDSs while two lines for promoter sequences (Fig 3).

**Fig 2.**
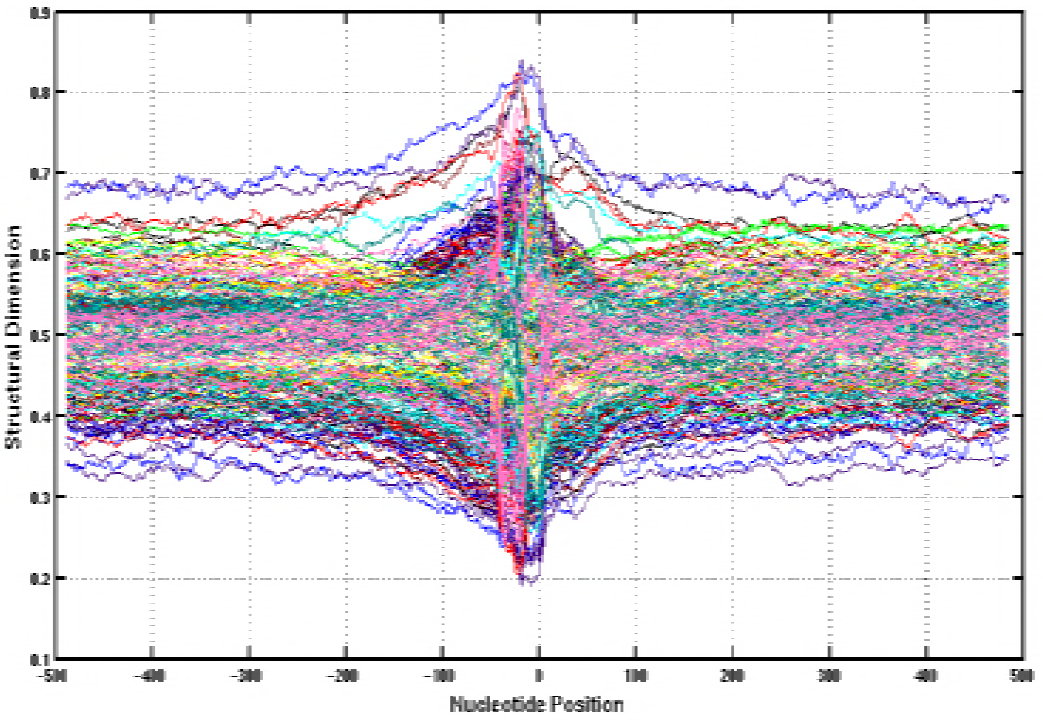
The normalized values of thirty one parameters (of all the twelve organisms) vs. nucleotide position with respect to TSS. Each organism was given single colour for all the 31 parameters. The plot represents 372 lines (31 × 12), a clear peak and cleft is observed at TSS or its adjoining upstream region.

**Fig 3.** The derived structural vectors profiles for the twelve organisms. Green lines represent sequences having TSS at “0”position while red line represents the CDSs. The green line showing a peak is vector1 while green line showing cleft is vector2; each obtained by combining normalized values of parameters showing same behaviour (see methods). For CDS, both vectors give a single line graph (red The ordinate represents the numeric value of the new structural scale while abscissa represents the nucleotide position relative to TSS

One surprising and striking observation was that despite their diversity, all organisms come to lie on the same position on this new structural scale (Fig. 3). The two vectors together give a uniform value of 0.5 for the CDSs of all organisms. For the promoter sequences, at TSS, vector1 gives a peak of magnitude ranging from 0.57 to 0.63 while vector2 yields a cleft of magnitude 0.3 to 0.37, for all the organisms. The above observation strongly indicates that DNA speaks a universal language.

### Different promoter sequences lead to similar structures

All the structural parameters act simultaneously to ultimately decide the DNA structure. The study was extended to generate structures of randomly selected promoter sequences, one from each organism (for sequence information, see supplementary Fig. S2). X3DNA software was used for generating structures using values of inter-Bp, intra-Bp and BP-axis, for the unique di-NT steps, generated during present study (Fig. 4).

**Figure 4:**
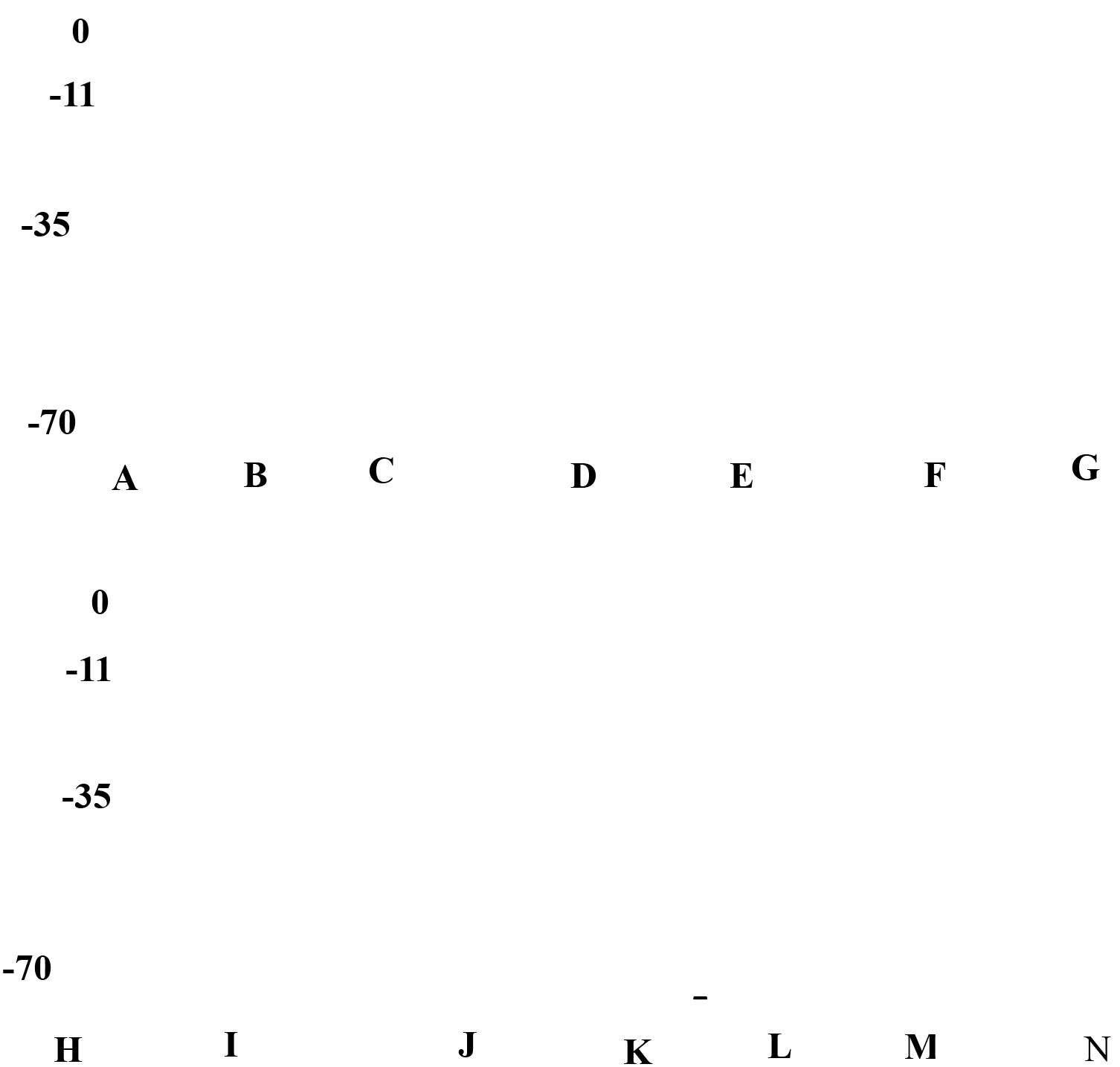
Three dimensional structures of promoter regions (−75 to +25 with respect to TSS) belonging to different organisms. A) *B. amyloliquefaciens*, (B) *C. pneumonae*, (C) *E. coli*, (D) *H. volcanii*, (E) *H. pylori*, (F) *M. psychrophilus*, (H) *M. tuberculosis*, (I) *P. aeruginosa*, (J) *S. typhimurium*, (K) *S. coelicolor, (*L) *S. species*, (M) *T. kodakarensis*. For comparison, similar structures of CDS region (g) and that of canonical B-DNA (n) are also given.

For comparison, structure of one CDS, randomly selected from *Bacillus amyloliquefaciens*, was also generated and a canonical B-DNA structure was also taken. All the promoter sequences despite poor sequence alignment (Supplementary Fig S2), led to almost similar structures, quite distinct from that of CDS and canonical B-DNA (Fig 4). This clearly indicates existence of a hidden structural code giving functional identities to promoter regions. As clear from Fig. 4, promoter regions adopt a slightly curved structure with variable groove dimensions throughout the length till TSS, on the other hand CDS and generic B-DNA adopt a straight structure with nearly uniform groove dimensions. Each promoter, however, displayed its own style of structural distortions, which might be unique to that organisms or that particular gene though much cannot be said at this stage.

To have a closer look on the structural changes in promoter region, 3D structures of five nucleotide long motifs from important regions (−11, −35, −70) of one promoter sequence (of *Bacillus amyloliquefaciens)*, were generated (Fig. 5).

**Figure 5:**
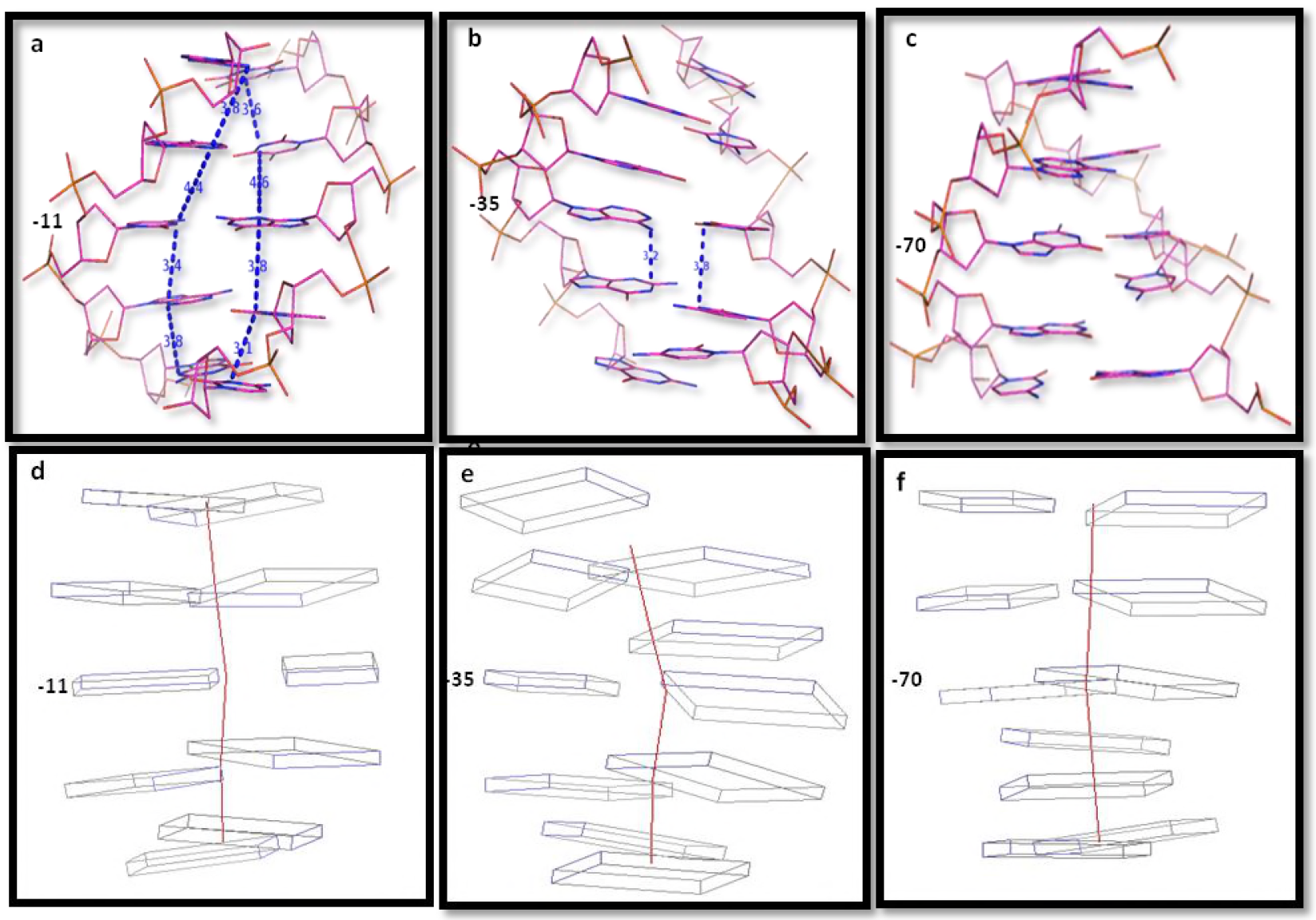
3D structures of 5 nucleotide long motifs of −11, −35 and −70 regions from one randomly selected promoter sequence of *Bacillus amyloliquefaciens*. a, b, c represent Line model structures of −11, −35 and −70 while d, e, f, represent their respective Calladine and Drew model structures.

As clear from Fig. 5, all the three regions show large deviations from the standard B-DNA in the arrangement of base pairs. At the −11 motif, a sharp increase in the vertical distance between base pairs at position −11 and −10 (4.4 & 4.6 Å in forward and reverse strand respectively) is distinctly visible; distance being significantly higher from 3.4 Å the standard inter-BP distance in B-DNA structure. This supports the observation made for the sharp increase in rise at the −11 region in Fig. 1. Another very interesting observation can be made from the Calladine model of this motif (Fig. 5d). The base pair at −11^th^ position shows high stretch but low twist, while consecutive base pairs on both sides exhibit high twist. Similar behaviour of twist (low twist position with high twist on both sides) and stretch for the −11 region was also recorded in Fig. 1.

The −35 motif displays remarkable deviations in the arrangement of base pairs (Fig. 5 b & e). The bp axis takes a slight bend at −35^th^ position. Base pairs at positions −34^th^ and −33^th^ show increased stagger, while all base pairs show variable level of tilt, roll, shift, slide, propeller twist. It is difficult to interpret conclusively from these structures but −35 motif definitely seems to be a hot spot of different structural deviations. For the −70 region, increase in angular distance from minor groove side (roll) is visible while a slight bending in axis is also observed (Fig. 5 c & f).

### Implications on transcription initiation

In the light of results discussed above, it seems that TSS and adjoining regions offer topographical signatures which act as strong nucleating factors for inviting RNAP and transcription factors. Topographical landscape of DNA molecular shapes have been considered to provide an efficient means of indirect readout of DNA (shape recognition) (32).

Further, the promoter structure and energetics seem to guide the subsequent interaction with various RNAP subunits and transcription factors. For instance, promoter DNA backbone undergoes a transition from BI to BII, resulting in placement of phosphate towards minor groove side; this might facilitate interaction with different domains of sigma subunit of RNAP e.g. α carbon terminal domains (αCTDs) of sigma (σ) subunit interacts with upstream promoter element using helix-hairpin-helix motifs (33) by hydrogen bonds between its backbone nitrogen and DNA backbone phosphate groups (34). Also, the −70 region and−35 region, important for interaction with various subunits of RNAP, show lots of structural deviations. What is the need for such visible and significant deviations in base pairs arrangements of these motifs? It demands a thorough investigation from many viewpoints. But at this stage it seems that these changes may provide a close access of its atoms to RNAP and other factors for required atomic interactions. Atomic details of −35 element recognition by σ4 of bacterial RNAP showed that helix-turn-helix motifs of σ4 interacts exclusively from major groove side on both template (35).

According to a recent report, promoter melting starts from within the −10 element (−12 to −7 nt position) by the interaction with σ_2_ subunit of RNAP resulting in flipping out of A_−11_ and T_−7_ bases of non-template strand which then get buried inside the pocket of σ_2_ subunit (36). Whether σ_2_ actively disrupts the base pairs-(A/T)_−11_ (T/A)_−7_ by its aromatic amino acids shovels or passively captures transiently exposed bases remains to be established (36,37). Present study offers some novel insights. At the−11 region, the vertical distance between two base pairs (rise) displays a sharp increase (Fig. 1), particularly between−11^th^ and −10^th^ position in one selected case (Fig. 5). This increased vertical distance between them might allow the aromatic amino acids shovels of σ_2_ to enter in the inter-base pair space. Such a significant increase in rise is not observed in other regions of promoter. Further, twist shows a typical behaviour: a sharp increase somewhere around−12^th^ position followed by sharp decrease and then again sharp increase (Fig. 1), the exact position of consecutive base pairs showing this pattern may vary from organism to organism but it is observed in all selected organisms (supplementary Fig S1-17). For the selected promoter of *Bacillus amyloliquefaciens*, it is−12,−11.−10 showing high, low and high twist respectively. It seems possible that under such conditions, the middle low twist position comes under strong torsional stain because of adjoining high twist regions and as a result either it gets partially extruded out or amino acid shovels of σ_2_ present in the inter-base pair space find it easier to extrude this unstable position base. Further investigations are needed to confirm this hypothesis. The energetics profile shows the −10 region to be the most unstable thus seems to facilitate promoter melting.

### Concluding remarks

Prevalent thinking posits that the RNAP is the key regulator of transcription initiation and after recognition and binding to the promoter DNA, it triggers a series of conformational changes in itself as well as in promoter DNA which are instrumental for transcription process initiation. However, the results obtained in the present study indicate that DNA exhibits changes in the overall structure at TSS and nearby regions without any aid from RNAP and transcription factors. Some previous studies also report that DNA dynamically directs its own transcription (38). On the basis of the results obtained in the present study, we conclude that DNA structure is a key regulator of transcription initiation; rather than acting as a passive platform on which RNAP acts to bring required changes, it assumes its structure and energetics on its own at TSS and nearby regions so as to offer conducive microenvironment to transcription machinery for precise recognition and atomic interactions needed for transcription initiation. Essentially, the message of TSS is already built into the structure and energetics of DNA sequences. Further, we have used the values for unique dinucleotide steps in our study. Since conformational, energetics and helical properties of a base pair are strongly influenced by nearest neighbours (18), we expect even better manifestation of these signals if tetra-nucleotide and higher order steps are considered instead of dinucleotides steps.

## Data Availability

We have considered 16519 primary promoter sequences and 6218 CDS sequence from 12 organisms (Table 1). User can download complete set or organism specific promoter sequence and CDS used in this analysis from our website (http://www.scfbio-iitd.res.in/software/data_TSS.jsp.). Rest of the data is available in the supplementary file.

## Supplementary Material

Supplementary material is available as separate file.

## Acknowledgements

Support from the Department of Biotechnology, Govt. of India to the Supercomputing Facility for Bioinformatics and Computational Biology (SCFBio), IIT Delhi, is gratefully acknowledged. The authors thank Professors Richard Lavery and Krystyna Zakrzewska for their helpful suggestions. PS extends thanks to Chaudhary Devi Lal University for granting Sabbatical Leave to her. AM is a recipient of UGC-SRF.

## Authors Contribution

BJ, PS, AM designed the project. PS, AM, PM collected the data. PS, AM, BJ analyzed the results and wrote the manuscript. MB, WO, KT, DB contributed some of the ideas and software used in the study and critically read the manuscript.

